# Catch bonds in sickle cell disease: shear-enhanced adhesion of red blood cells to laminin

**DOI:** 10.1101/2022.11.12.515898

**Authors:** Utku Goreke, Shamreen Iram, Gundeep Singh, Sergio Domínguez-Medina, Yuncheng Man, Allison Bode, Ran An, Jane A. Little, Christopher L. Wirth, Michael Hinczewski, Umut A. Gurkan

## Abstract

Could the phenomenon of catch bonding—force-strengthened cellular adhesion—play a role in sickle cell disease, where abnormal red blood cell (RBC) adhesion obstructs blood flow? Here we investigate the dynamics of sickle RBCs adhering to a surface functionalized with the protein laminin (a component of the extracellular matrix around blood vessels) under physiologically relevant micro-scale flow. First, using total internal reflectance microscopy we characterize the spatial fluctuations of the RBC membrane above the laminin surface before detachment. The complex dynamics we observe suggest the possibility of catch bonding, where the mean detachment time of the cell from the surface initially increases to a maximum and then decreases as a function of shear force. We next conduct a series of shear-induced detachment experiments on blood samples from 25 sickle cell disease patients, quantifying the number and duration of adhered cells under both sudden force jumps and linear force ramps. The experiments reveal that a subset of patients does indeed exhibit catch bonding. By fitting the data to a theoretical model of the bond dynamics, we can extract the mean bond lifetime versus force for each patient. The results show a striking heterogeneity among patients, both in terms of the qualitative behavior (whether or not there is catch bonding) and in the magnitudes of the lifetimes. Patients with large bond lifetimes at physiological forces are more likely to have certain adverse clinical features, like a diagnosis of pulmonary arterial hypertension and intracardiac shunts. By introducing an *in vitro* platform for fully characterizing RBC-laminin adhesion dynamics, our approach could contribute to the development of patient-specific anti-adhesive therapies for sickle cell disease. The experimental setup is also easily generalizable to studying adhesion dynamics in other cell types, for example leukocytes or cancer cells, and can incorporate disease-relevant environmental conditions like oxygen deprivation.

**SIGNIFICANCE:** Abnormal red blood cell adhesion to the walls of blood vessels is a central feature of sickle cell disease. We study this adhesion by experimentally measuring how long on average red blood cells adhere to a protein-covered surface, and how the strength of the cell-protein bond depends on the force resulting from the surrounding fluid flow. The results vary widely from patient to patient, with some cases showing an unusual regime where the mean bond strength increases with force. We connect these measurements to clinical aspects of the disease, which may aid in the design of individualized therapies in the future.

## INTRODUCTION

Red blood cells (RBCs) are packed with oxygen-binding hemoglobin molecules. In sickle cell disease (SCD), sickle hemoglobin polymerizes when deoxygenated due to a mutation of a single gene (1). Polymerized sickle hemoglobin induces physical and chemical changes (2, 3) that make sickle RBCs (sRBCs) stiffer (4–6), more dense (7) and abnormally adhesive (8–10) relative to healthy RBCs. These changes in biophysical properties of sRBCs may initiate a cascade of events leading to a disease complication known as a vaso-occlusive crisis, where the sRBCs aggregate and block blood vessels (11, 12). Part of the molecular mechanism underlying the crisis is enhanced sRBC adhesion to extracellular matrix proteins, which constitute the scaffold of blood vessels. In sickle cell disease, extracellular matrix proteins may be exposed to the bloodstream due to elevated levels of circulating endothelial cells that have been shed away from the walls of the blood vessels (13). These levels spike dramatically at the onset of a vaso-occlusive crisis (14). Among extracellular matrix proteins, laminin (LN) mediates the most avid sRBC adhesion (8, 15). In this work we study how the abnormal adhesion of sRBCs to LN depends on the shear force that arises from the surrounding fluid flow.

Monotonic decrease of a bond’s lifetime with increased force is a characteristic of slip bonds (16), and is the default expectation for most biological adhesion interactions. In contrast, we find that sRBC-LN adhesion for a subset of patients with SCD exhibits a non-monotonic lifetime response to shear, *i.e*. having a regime where adhesion strength seems to counter-intuitively rise with increasing shear force (Fig. 1). This behavior is known as catch bonding (17–20), and allows the system to potentially withstand wider ranges of physiological forces. Though catch bonding is often considered at the level of individual protein-ligand complexes, in our study we focus on the larger scale phenomenon of an entire sRBC cell detaching from a protein-functionalized surface. Here the effective “bond” lifetime is the collective result of many individual interactions between surface proteins and membrane ligands. This complexity opens the door for non-trivial adhesion dynamics. In fact, as discussed below, the existence of catch bonds necessarily requires complex detachment dynamics, for example multiple detachment pathways whose likelihood varies a function of force (21). Catch bonds are not new for RBCs—they were previously observed in cytoadhesion of *P. falciparum-infected* RBCs in the placental intervillous space (22). However the present study is the first to find catch bond behavior in sRBC-LN adhesion, and to try to understand its implications in the context of sickle cell disease.

**Figure 1:**
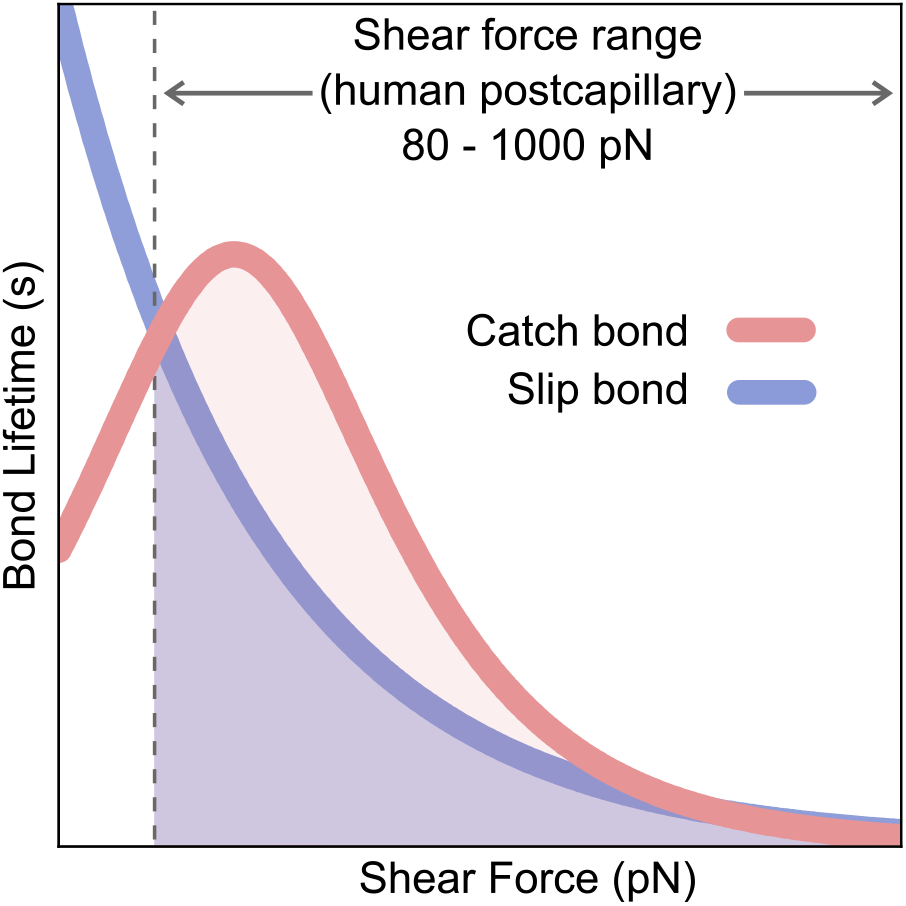
Illustration of slip versus catch bond behaviors for an adhered sRBC under shear flow. Increased shear force could either shorten the bond lifetime monotonically (slip bond) or initially enhance the lifetime and strengthen the adhesion (catch bond), before reducing it again at larger forces.

Although the molecular basis and biochemical mechanisms of sRBC-LN adhesion are well characterized (13, 23–25), the hemodynamical aspects of this adhesion are only partially understood. The sRBC-LN adhesion literature discusses topics such as RBC deformation during detachment (26) and the number of adhered RBCs under different shear conditions (27, 28). But key questions on how forces due to blood flow regulate the bond dynamics, and the heterogeneity of adhesion characteristics between different patients, remain open. Though SCD stems from a single point mutation (3), intriguingly there is great population level phenotypic heterogeneity, probably due to genetic and epigenetic factors, age, and geographical origins (29, 30). Additionally, heterogeneity is also observed at the level of biochemical signatures and morphology among cells of the same SCD patient (31, 32). Thus it is natural to ask whether heterogeneity also exists in the dynamics of sRBC-LN adhesion, and whether it could be linked to larger-scale phenotypic heterogeneities among SCD patients.

To comprehensively investigate sRBC-LN adhesion, we have developed a multi-faceted experimental framework, supported by theoretical modeling of the bond dynamics. First, to get a detailed look at the sRBC detachment process from an LN-functionalized surface, we report real-time, label-free probing of individual single-cell detachment events using Total Internal Reflectance Microscopy (TIRM). To the best of our knowledge, this is the first time TIRM has been applied to an RBC. Next, we conduct experiments in LN-functionalized microchannels, observing sRBC detachment under both sudden force jumps and linear force ramps. Using a model for the detachment dynamics under a force ramp, we can extract key dynamical characteristics—including the mean lifetime versus force profile that indicates the presence or absence of catch bonds. Finally, we examine how these characteristics vary among a cohort of SCD patients, and check whether they correlate with clinical observables.

## RESULTS AND DISCUSSIONS

### Single-cell detachment under TIRM: dynamic variation of adhesion sites

Polymerization of sickle hemoglobin takes place rapidly after the formation of nuclei (33). This rapid expansion of stiff and rod-like sickle hemoglobin polymer causes flip-flop of membrane lipids and proteins in clusters and these clusters become adhesion sites (3, 34, 35). Our first research question focuses on the dynamics of these adhesion sites during the detachment of an sRBC. Previously, adhesion sites on RBCs were visualized with confocal microscopy and total internal reflectance fluorescence microscopy (35, 36). The time scale of adhesion site switching is likely to be much smaller than the acquisition times often needed for fluorescence imaging that is required for these two methods. Also, both of these methods involve antibody staining that could potentially interfere with the ligands. TIRM is an existing method for label-free, high temporal resolution investigation of spherical particle-surface interactions. Our team is developing a new class of TIRM, named Scattering Morphology Resolved TIRM (SMR-TIRM), suitable for probing interactions between anisotropic particles and a neighboring surface. We recently described the implementation of SMR-TIRM on anisotropic particles (37, 38). This improvement expands the applicability of TIRM to biological targets.

We mapped the spatial proximity between an sRBC and the LN functionalized surface qualitatively during detachment under a gradually increasing shear force with our refinement of TIRM. Fig. 2 shows a representative result for a single sRBC. To quantify the actual closeness of the interfaces, total internal reflectance (TIR) as a function of orientation was needed. Therefore, we normalized the intensity traces arbitrarily to the average intensity before detachment, in which the sRBC was mostly static on the surface. TIR intensity (a.u.) the correlates with the normalized proximity between the sRBC and the LN surface. Also, the integrated TIR intensity indicated the averaged proximity over the whole cell surface. We assumed that, probabilistically, the closer the distance between the interfaces the greater the likelihood of adhered ligands. We observed a few significant phenomena during the detachment and illustrated these in Fig. 2 with timestamps (t_i_) that specify the distinct phases of detachment. The first phenomenon was that the TIR intensity varied spatially over the surface of the sRBC during the progression of detachment. The TIR intensity increased around the leading edge between t_1_ and t_2_ (Fig. 2). Also, the high TIR intensity region at the trailing edge shifted in the traverse direction between t_1_ and t_2_. The second phenomenon was that during the second phase of detachment the integrated TIR intensity increased to 1, relative to its initial value of of 0.75. It should be noted that at t_3_, the sRBC was released from the boundary (LN surface) and the integrated TIR intensity during this phase dropped to approximately 0.5. This kind of intensity change is roughly consistent with the average distance between sRBC and the LN surface increasing on the order of 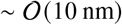 from the initial equilibrium distance, although a more precise measure of height change would require a deeper understanding of how scattering depends on the orientation and shape of the RBC (37, 38). The sRBC displayed a dragging motion until t_4_ and tumbled at t_5_. The spike of integrated TIR intensity at t_5_ was considered an artifact because turning or flipping of the RBC after detachment may also contribute to the overall scattering intensity, as the cross-section of the cell changes with respect to the propagation vector of the evanescent wave. The video of the TIRM experiment illustrated in Fig. 2 can be accessed as Supplementary Video 1. The details of the shear force calculations can be found in the Supplementary Information (SI).

**Figure 2:**
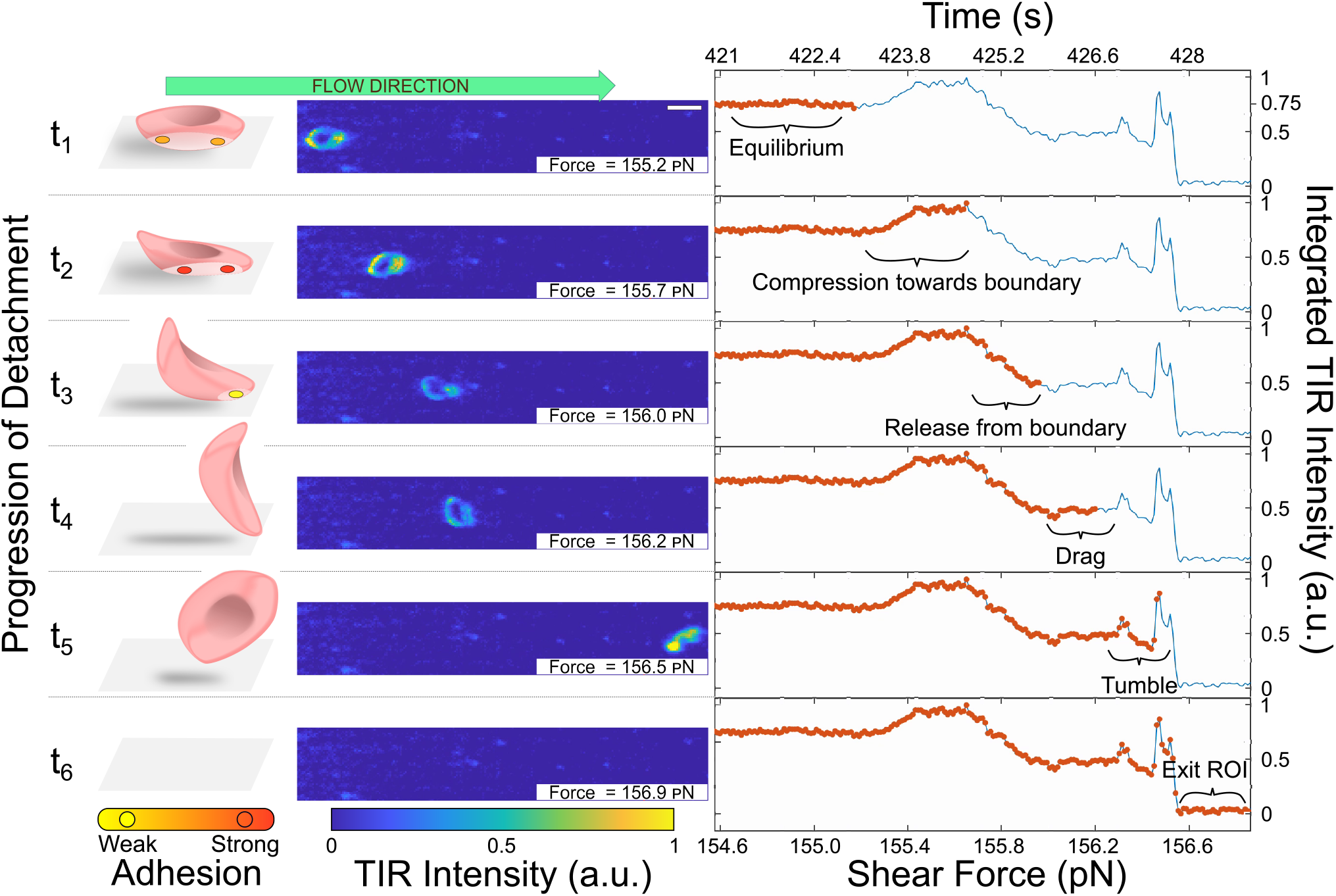
The proximity between a detaching red blood cell and laminin surface is dynamically resolved by Total Internal Reflectance (TIR) microscopy under a microfluidic physiological flow ramp. The distance between the cell and the laminin surface (boundary) is inversely correlated with TIR intensities. The smaller this distance, the greater the likelihood of binding. The scattering intensity landscape shows an initial decrease in distance at both the leading and trailing edges of the cell (timepoints t_1_ to t_2_). This decrease can also be inferred from integrated TIR which increases from 0.75 to 1. Subsequently (t_2_ to t_3_) this value decreases to 0.5 as the cell is released from the boundary, before eventually tumbling, detaching, and exiting the region of interest (ROI). Dynamic variations in cell-surface proximity during the detachment process under gradually increasing shear force indicate a complex underlying binding scenario.

With TIRM, the detachment of sRBCs was observed as an elevation of the scattered light intensity, followed by a decrease and brief moments of post-detachment contact seen as peaks in the intensity traces. Furthermore, the change in integrated TIR intensity was complemented by spatial variations of TIR intensity. Though we show the results for one sRBC, similar variations are generic features across different cell observations. TIRM reveals that detachment process under changing shear forces likely involves a dynamic interplay of different adhesion sites along the cell membrane, as well as varying distances between the membrane and surface at any given site. The collective contribution of these microscopic phenomena means there is the possibility of having complex overall detachment dynamics. In Ref. (21) it was argued mathematically that for the multidimensional free energy landscape of a bond, there are two scenarios that could lead to non-monotonic bond lifetime vs. force: i) multiple physically distinct detachment pathways, as could happen in our case if detachment occurs stochastically through different combinations of attachment sites; ii) a detachment pathway where bond extension initially decreases with increasing force, before increasing at a later stage—compatible with our observation of initial compression and then release between t_1_ and t_3_. While neither of these scenarios means that a catch bond must necessarily exist in the system, having one of them (or a combination of both) is a prerequisite for catch bonding. Thus our TIRM results indicate that sRBC-LN adhesion certainly has the potential for behaving like a catch bond. To determine whether this potential is realized, we turn to force jump and force ramp microfluidic experiments.

### Force-strengthened sickle red blood cell adhesion

As a first step toward identifying possible catch bond behavior, we measured net RBC adhesion on an LN-functionalized surface following a sudden jump in shear force. We used four different jump magnitudes (Fig. 3a): in each case the system was equilibrated at an initial shear force of 50 pN for 15 minutes, and then either kept at the same force (as a control), or stepped up to 150, 250, or 450 pN. The force values all fall within the physiological range observed in post-capillary venules. We monitored the same 32 mm^2^ area of the surface, capturing images at two minute intervals. To obtain the net RBC adhesion per *μ*L of blood sample, we take the mean difference of the number of adhered RBCs between consecutive imaging intervals after the jump and divide it by the sample volume that has passed through the microchannel during each interval. The resulting net RBC adhesion values are shown in Fig. 3b for samples from seven patients, plotted versus the shear force after the jump. The net adhesion at lower forces is positive, reflecting the fact that on average more cells attach than detach during every imaging interval under those conditions. However the way that adhesion depends on force varies dramatically among different patients. Four out of the seven patients exhibit clear indications of a force-strengthened (“catch-like”) adhesion: a significant increase in net adhesion for the 150 pN jump compared to the 50 pN control case. At higher forces the net adhesion steadily decreases. For the remaining three patients the peak in net adhesion versus force was either marginally discernible, given the error bars, or not present (“slip-like”).

**Figure 3:**
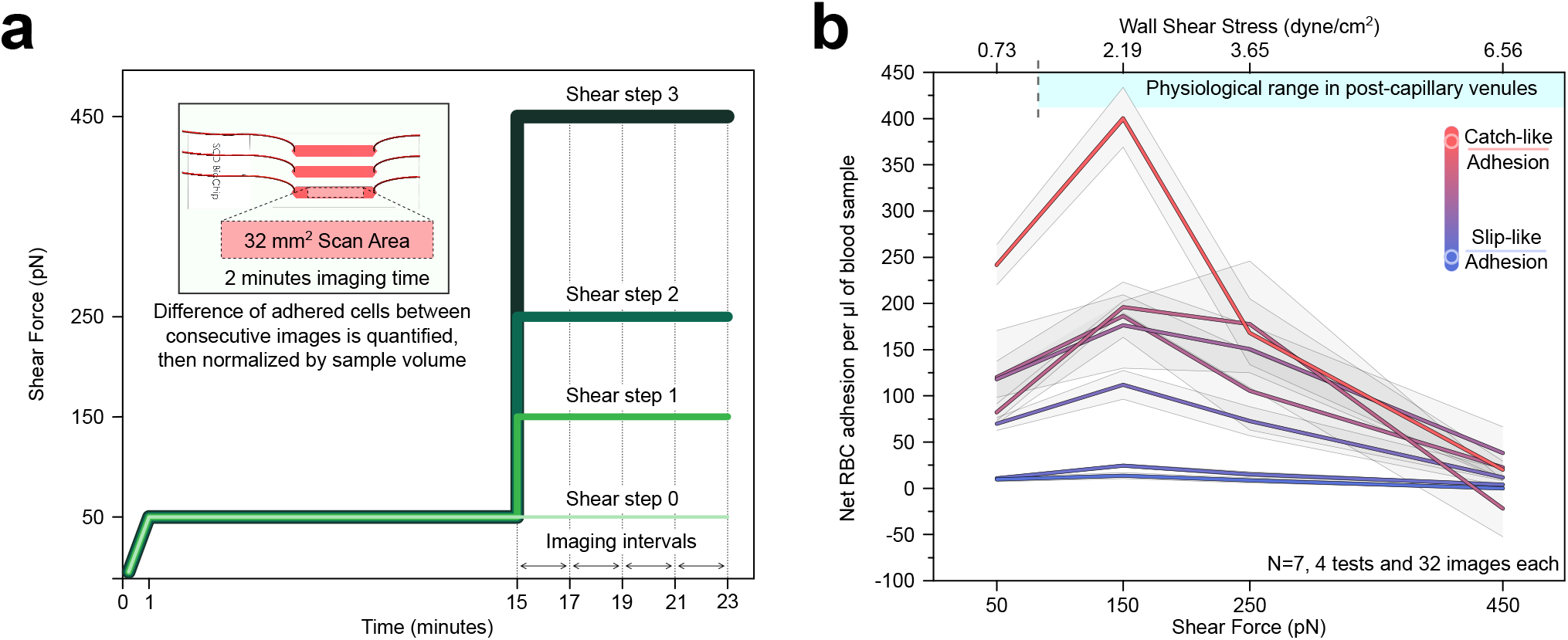
Net sRBC adhesion normalized by the sample volume follows a heterogeneous force-strengthening pattern in the physiological range of shear forces of post-capillary venules. **(a)** Shear force profiles for the force jump experiments and an illustration of the microfluidic device (inset). Consecutive microscopic images are taken at two minutes intervals and then the change in the number of adhered sickle RBCs is normalized by the sample volume that resulted in that change. **(b)** Net RBC adhesion after the force jump as a function of the final force value for seven homozygous sickle cell disease patients. Net adhesion curves vary widely between patients, with some showing clear “catch-like” strengthening in the 50-150 pN regime and others exhibiting more “slip-like” characteristics. Error bars represent the standard error of the mean.

While force-enhanced net adhesion among the first subset of patients is suggestive of catch bonding, this experimental setup does not directly measure bond lifetime versus force. In order to get a more fine-grained picture of sRBC-LN bond dynamics, we need to not just count total numbers of cells on the surface at any given time, but track individual adhered cells until they detach. This is what we implemented in the next series of experiments.

### Shear ramp experiments: extracting mean bond lifetime versus force

The most direct probe of catch vs. slip bonding would be measuring the mean bond lifetime *τ*(*f*) as a function of a constant applied force *f*, as shown qualitatively in Fig. 1. In principle this can be implemented by repeatedly subjecting individual adhered cells to a particular force value *f* until detachment occurs (for example using atomic force microscopy (39)), collecting sufficient statistics, and then repeating this process for a range of different *f* values. However, when considering the attachment of an entire sRBC cell to the LN surface, mediated by multiple adhesion sites, the bond lifetime at low shear forces can be quite long (minutes or tens of minutes). Hence collecting sufficient statistics to reliably measure *τ*(*f*) would be prohibitively time consuming, particularly if the goal is to repeat the analysis across many different SCD patient samples. The more practical alternative in this case is to study detachment dynamics in an LN-functionalized microfluidic channel (27, 28, 40). By linearly increasing shear force in the channel with time at some constant rate *r*, we can ensure that eventually most of the initially adhered sRBCs will detach, yielding a large number of observed detachment events. However to convert these raw observations into an estimate for *τ*(*f*) requires several steps of data analysis and modeling, which we summarize below.

The first step is to estimate the survival probability density Σ*_r_* (*f*) for the adhered sRBCs at ramp rate *r*, where Σ*_r_* (*f*) is the probability that a cell adhered at the start of the force ramp is still adhered when the force has reached a value *f*. To do this, we need to be able to track individual adhered sRBCs, measure the time of detachment, and hence obtain the corresponding force at detachment. A well populated, large field of view of preset size is chosen at the start of the assay and the initial number of adhered sRBCs is noted. The field of view is locked in for the duration of the assay, and videos with a top-down view are recorded. The videos are analyzed using a custom automated image processing pipeline, building on a deep neural network approach we recently developed (41). We can find the survival function via Σ*_r_*(*f*) ≈ *N*(*f*/*r*)/*N*(0), where *N*(*t*) the number of adhered sRBCs left at time *t* after the start of the ramp. Fig. 4a shows the four different force protocols used in the assay, with varying ramp rates. The corresponding estimates of Σ_*r*_(*f*) for one patient are shown in Fig. 4b. The initial number of adhered cells *N*(0) within our field of view varies between experimental trials and different patients, but is of order 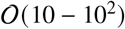 cells.

**Figure 4:**
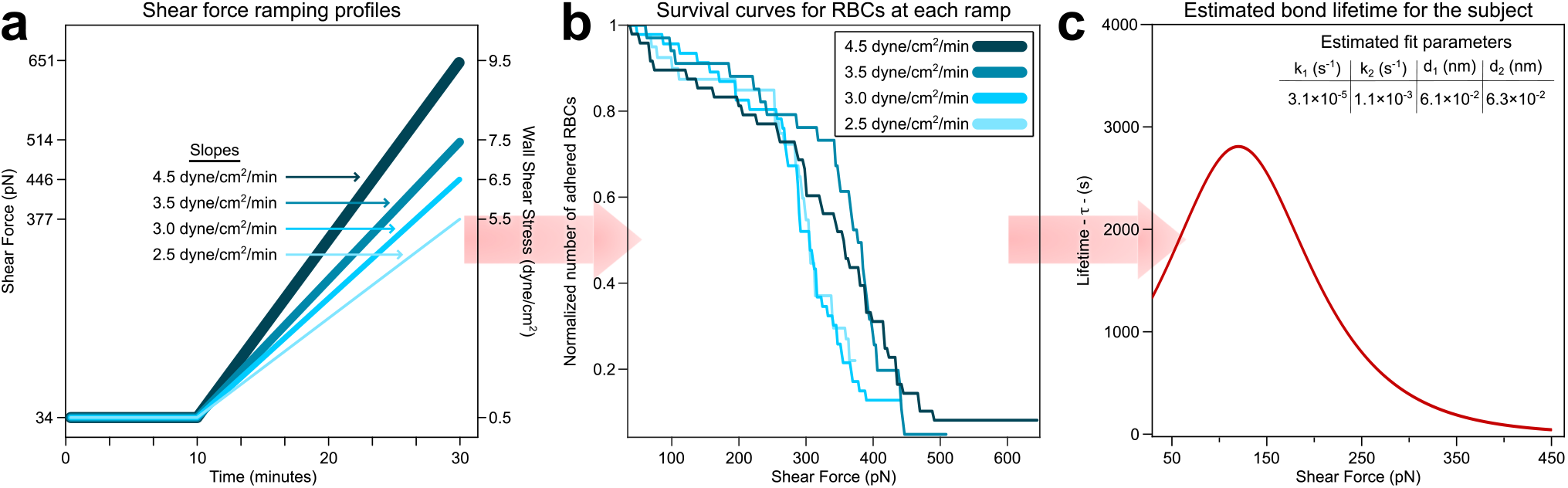
Experimentally obtained survival curves under varying shear force ramping yield an estimate for the mean bond lifetime as a function of force. **(a)** Shear force profiles for the ramped shear experiments, with four different ramp rates *r*. **(b)** Survival curves Σ_*r*_(*f*) for sRBC-LN adhesion under the ramps shown in panel a, based on blood samples from one SCD patient. **(c)** Using the Σ_r_(*f*) data, we employ maximum likelihood estimation to fit the parameters of the model in Eqs. (1)–(2). The four fitted parameters (*k*_1_, *d*_1_, *k*_2_, *d*_2_) allow us to predict the mean bond lifetime *τ*(*f*) versus force *f* (red curve) for this patient.

The second step is to relate the *Σ_r_*(*f*) estimates to the mean bond lifetime *τ*(*f*) at constant force. There is one case where this relationship is easy to formulate: the pure adiabatic pulling regime (42). To define this regime, let us assume our system is characterized by heterogeneous bound states that can dynamically interconvert between each other before detachment—as is suggested by the TIRM results, and typical for catch bonds (43). For a given ramp rate *r*, we are in the pure adiabatic pulling regime if the timescale of interconversion, as well as the timescale of equilibration within each state, are both much much smaller than the typical timescale of detachment at that value of *r*. In other words, the system has a chance to extensively sample the bound states before detachment, and the overall bond lifetime reflects an averaging over those states. If we are in this regime, *Σ_r_*(*f*) and *τ*(*f*) are related via (42, 44):

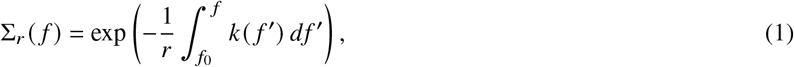

where *k*(*f*) ≡ 1/*τ*(*f*) is the mean detachment rate at force *f*, and *f*_0_ is the force at the beginning of the ramp (*f*_0_ ≈ 34 pN in our case). As can be seen from the structure of Eq. (1), there is a straightforward empirical test for whether the adiabatic regime is actually realized (42): the quantity −*r* log Σ*_r_*(*f*) should be independent of *r* across the range of measured forces (within experimental error bars). We can readily verify this criterion using our estimates of *Σ_r_*(*f*) at four different ramp rates *r*, and we find that for our experimental setup the pure adiabatic regime holds (see SI Figure 1).

In principle Eq. (1) can be inverted to find *k*(*f*) = −*r*(*d*/*df*) ln Σ*_r_*(*f*), and hence *τ*(*f*) = 1/*k*(*f*), but taking derivatives of noisy estimates of Σ*_r_*(*f*) is typically unreliable. Hence we adopt an alternative approach, where we assume an underlying functional form for *k*(*f*) and use maximum likelihood estimation (on the combined data from all four ramp rates, for better statistics) in order to find the parameters of the *k*(*f*) function. The full details of the parameter estimation procedure are in the SI. The form of *k*(*f*) should be flexible enough to cover both slip and catch bonds behaviors, and a reasonable starting point is the phenomenological model of Ref. (43). This model has two bound states with distinct detachment pathways, but when the timescales of interconversion between bound states are fast (our regime of interest), the detachment rate reduces to a simple two-exponential form (43, 45):

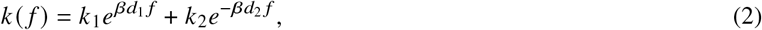

where *β* = (*k_B_T*)^-1^, *T* is temperature (taken to be *T* = 298 K), and *k_B_* is the Boltzmann constant. The first term has the same structure as the Bell model for slip bonds (16), with a rate coefficient *k*_1_ and a parameter *d*_1_ (measured in units of distance) that characterizes the sensitivity of the Bell term to force *f*: “brittle” (less force-sensitive) bonds correspond to smaller *d*_1_, and “ductile” (more force-sensitive) bonds to larger *d*_1_ (46). The second term has a similar form, with parameters *k*_2_ and *d*_2_, but the sign difference in the exponential means this term can potentially increase the overall rate (and hence decrease lifetime) at small forces, leading to catch bond behavior. The latter typically requires the second term to be dominant at small forces, with *k*_2_ ≫ *k*_1_. If instead the first term dominates, we end up with a slip bond. Fig. 4c shows the parameter estimation results for the survival data in Fig. 4b, with the corresponding *τ*(*f*) curve for that particular patient. The curve turns out to exhibit clear catch bond behavior, but how common is this among SCD patients in general? To investigate the variability of *τ*(*f*), we turn to analyzing a cohort of 25 SCD patients.

### Variation in bond dynamics among a group of SCD patients

Fig. 5a,b summarizes the fitting results for the 25 SCD patient samples, with Fig. 5a showing the (*k*_1_, *d*_1_) parameters of the Bell term in Eq. (2), and Fig. 5b showing the (*k*_2_, *d*_2_) parameters of the catch term. 9 out of the 25 patients had non-negligible *k*_2_ values, and hence exhibited catch bond behavior (or near-catch-bond behavior, in the case where *τ*(*f*) levels off at small forces but does not reach a peak). This is seen clearly in Fig. 5c, which plots *τ*(*f*) for all the patients (catch-like curves in red, slip-like curves in blue). The (*k*_1_, *d*_1_) parameters in Fig. 5a obey a roughly exponential relationship (dotted line), with smaller *k*_1_ correlated with larger *d*_1_. Explaining this curious relationship (and whether it is a universal feature of SCD patient populations) is an open question for future studies. An answer likely requires going beyond the phenomenological theory of Eq. (2) to more detailed, structure-based models of the various adhesion interactions that collectively contribute to sRBC-LN bond dynamics (along the lines of Refs. (47, 48)).

**Figure 5:**
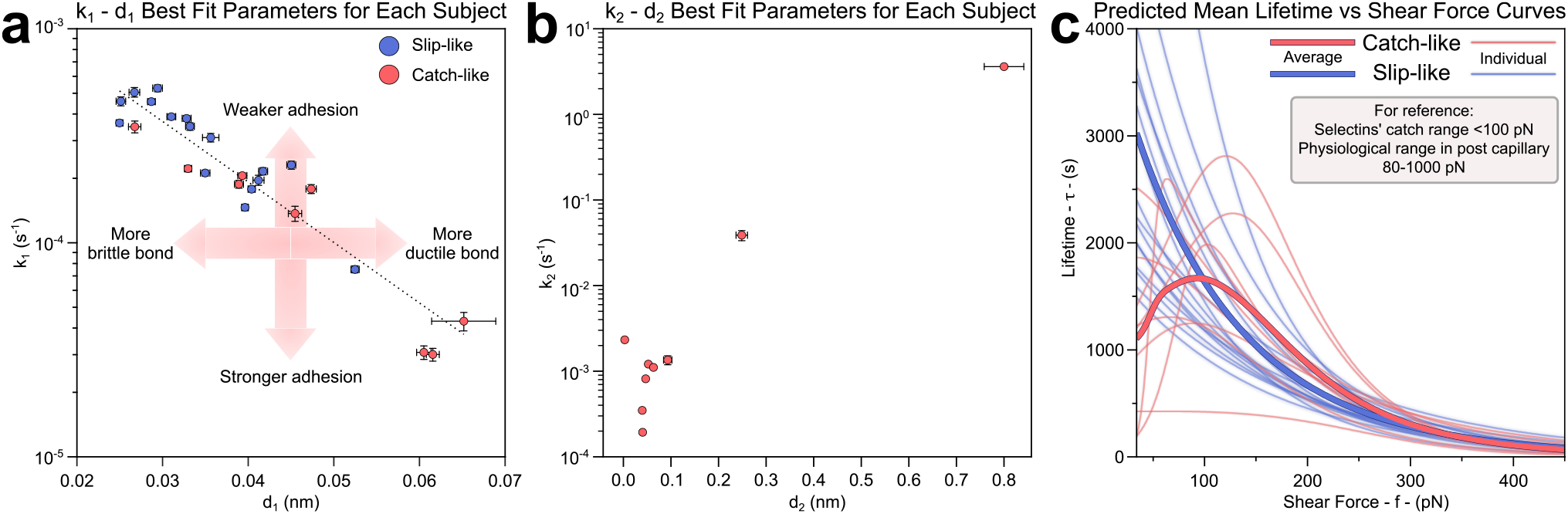
Bond lifetime fitting results for a group of 25 SCD patients. **(a)** (*k*_1_,*d*_1_) best-fit values, characterizing the first (Bell) term in Eq. (2). The parameters exhibit an approximately exponential trend (dashed line). **(b)** (*k*_2_, *d*_2_) best-fit values, corresponding to the second (catch) term in Eq. (2). Only 9 out of the 25 patients are shown here, the ones with non-negligible *k*_2_ values that lead to catch-like bond behavior. The same 9 patients are highlighted in red in in panel a. **(c)** Best-fit predictions of the mean bond lifetime *τ*(*f*) for individual patients are plotted as thin curves. Slip-like and catch-like behavior is colored in blue and red respectively. The average lifetime behaviors for these two subgroups are plotted as thick curves.

The variation in *τ*(*f*) among the 25 patients in Fig. 5 is quite dramatic, particularly at smaller forces. Near the lower end of the force range typical of post-capillary venules (~ 80 pN), the mean adhesion times vary from hundreds to several thousand seconds. For those cases that do show catch bonding, the locations of the peaks (~ 50 – 150 pN) are similar to peaks observed in Fig. 3b, corroborating the force jump experimental results. The *τ*(*f*) curves averaged over all the catch-like and slip-like patients are shown as thick red and blue curves respectively. These subgroup results indicate that on average catch bonds play a mechanical buffering role, increasing the bond lifetime above the slip subgroup for physiologically relevant forces (≳ 100 pN). (Though it is worth noting there are individual slip bond cases with comparable or larger lifetimes to the catch bond cases.) Does such strengthened adhesion play a role in the clinical manifestations of SCD? We tackle this question in the next section.

### Bond lifetime correlates with clinical features of SCD

Catch bonds can be useful physiologically—for example, they are involved in selectin-mediated rolling of leukocytes, a mechanism that helps leukocytes recognize the site of inflammation in blood vessels (49). In this instance, catch bonds act as a sensor of increased shear force due to vasoconstriction, helping to regulate the desired function of the cells. Catch-like sRBC adhesion in SCD could use the same sensor mechanism to deleterious effect, leading to adhesion of sRBCs around sites of inflammation. For comparison, the peaks in bond lifetimes for selectins with their ligands occur in a similar force regime (<100 pN(50)) to the sRBC-LN catch bonds. Thus, catch-like sRBC adhesion may be detrimental for individuals with SCD who suffer from chronic inflammation and even though the anti-adhesive crizanlizumab inhibits leukocyte adhesion, the shear-enhanced adhesion of sRBCs may still persist. Another instance of a pathology that causes elevation of shear force is hyperperfusion—increase of blood velocity. Hyperperfusion is both a systemic and regional dysregulation in SCD (51, 52), and contributes to the perfusion paradox that instigates vaso-occlusion in postcapillary venules (53). The combination of sRBC catch-like adhesion and hyperperfusion could potentially magnify adverse outcomes.

To test whether the observation of catch-like behavior by itself has SCD clinical associations, we conducted a statistical analysis of the catch-like versus slip-like subgroups in our cohort of 25 SCD patients. Examining a variety of clinical patient parameters related to SCD phenotypes, we found no significant clinical associations between the two groups (Supplementary Table 1). This is likely due to the fact that both catch-like and slip-like patients exhibit wide variation in their *τ*(*f*) behaviors, as discussed in the previous section. For example at physiological forces there are both catch-like and slip-like patients among the ones with highest bond lifetimes (and also among the lowest).

Given this fact, we suspected that a better observable might be the bond lifetime under force, since larger adhesion lifetimes should have clear biological consequences in the context of the disease. For each patient, we averaged the *τ*(*f*) result from Fig. 5 over the physiological force range accessible to our measurements, 80-450 pN. This defines a force-averaged bond lifetime 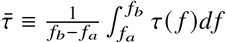, where *f_a_* = 80 pN, *f_b_*, = 450 pN. The distribution of 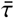 values for our 25 patients was normal but slightly skewed, and we divided the group into two using the median rank (Fig. 6A, p=0.102, Anderson-Darling test, skewness=0.67). Supplementary Fig. 2 replots the *τ*(*f*) results of Fig. 5, but now highlighting the two groups based on low and high 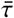. Strikingly, the SCD phenotypes that are related to inflammation and elevated blood velocity were more pronounced in the high 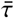 group than in the low 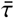 group (Table 1). These differences are also shown as a distribution in Fig. 6b, c: 10.9±3.1 white blood cells (in units of 10^9^/L) in the low 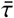 group vs. 13.4±2.1 in the high 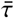 group (*p* = 0.032); similarly, 2.39±0.43 m/s tricuspid regurgitation velocity (TRV) for low 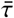 vs. 2.90±0.52 m/s for high 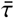 (p=0.011, one-way ANOVA). It is important to note that, as we have discussed above, both of these phenotypes manifest themselves with a regional or systemic increase of shear force.

**Figure 6:**
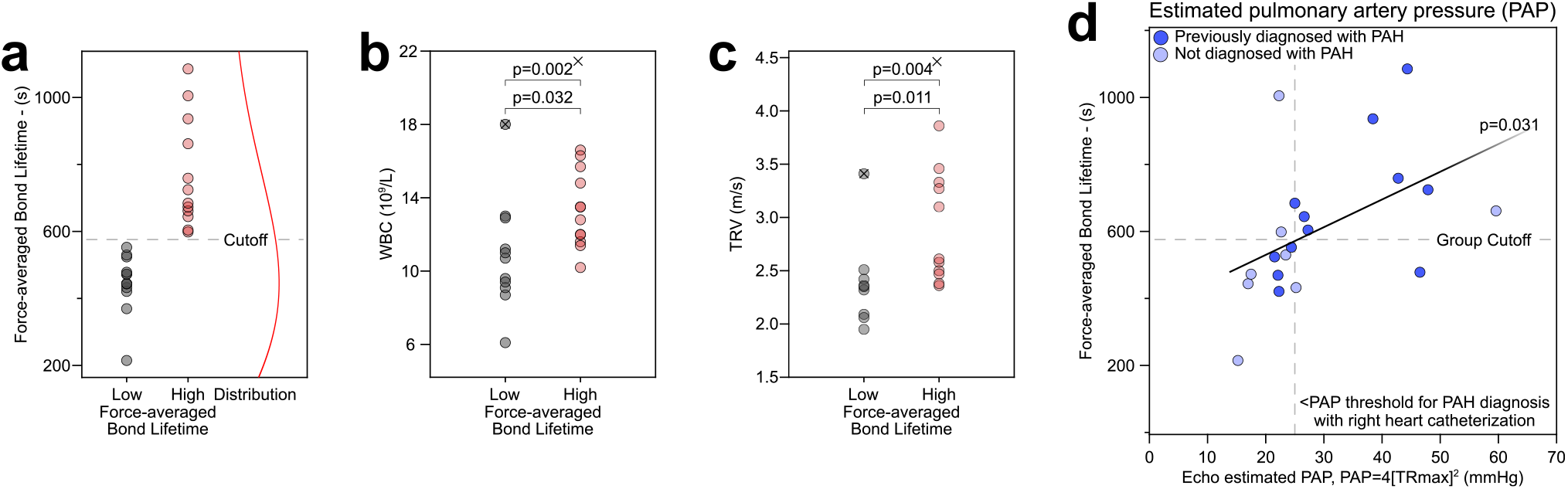
High force-averaged bond lifetime 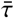 is associated with clinical markers that suggest the presence of a regional or systemic increase in shear force. **(a)** distribution of 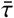 values among 25 SCD patients, and the cutoff for dividing the group into high and low 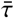 subgroups. **(b)** White blood cell (WBC) count and **(c)** tricuspid regurgitation velocity (TRV) are associated with high 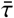. The × signs denote the outliers (*p* = 0.035 & *p* = 0.003 for WBC and TRV, respectively, Dixon’s Q ratio test) and the *p* values with × superscripts denote the statistical significance in the absence of the outliers. **(d)** 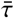 also correlates with echocardiogram-estimated pulmonary artery pressure (PAP). Echocardiograms are important for the diagnosis of pulmonary artery hypertension with right heart catheterization.

**Table 1:** Clinical phenotypes of 25 SCD patients based on force-averaged bond lifetime 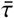. High 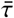 is associated with larger white blood cell counts and peak tricuspid regurgitation velocities, as well as diagnoses of pulmonary artery hypertension and intracardiac shunts.

| Parameters (Units) | Subjects with<br>low force-averaged<br>bond lifetime<br>N=12 | Subjects with<br>high force-averaged<br>bond lifetime<br>N=13 | p |
| --- | --- | --- | --- |
| Age | 33.2 ± 6.2 | 33.2 ± 9.5 | 0.997* |
| **White Blood Cells (10<sup>9</sup>/L)** | **10.9 ± 3.1** | **13.4 ± 2.1** | **0.032\*** |
| Absolute Neutrophil Count (10<sup>6</sup>/L) | 6630 ± 3169 | 8454 ± 1657 | 0.098* |
| Platelets (10<sup>9</sup>/L) | 369.7 ± 62.0 | 355.1 ± 127.6 | 0.734* |
| Mean Corpuscular Volume (fL) | 86.5 ± 15.9 | 89.3 ± 3.6 | 1.000** |
| Hemoglobin (g/dL) | 8.3 ± 1.5 | 8.9 ± 1.5 | 0.346* |
| Hematocrit (%) | 24.6 ± 2.9 | 25.9 ± 4.7 | 0.449* |
| **Tricuspid Regurgitation Velocity, Peak (m/s)** | **2.39 ± 0.43** | **2.90 ± 0.52** | **0.011\*\*** |
| Red Blood Cells (10<sup>12</sup>/L) | 2.7 ± 0.33 | 2.9 ± 0.6 | 0.305* |
| Absolute Reticulocyte Count (10<sup>9</sup>/L) | 412 ± 183 | 487 ± 170 | 0.317* |
| Reticulocyte percent (%) | 15.0 ± 5.7 | 17.6 ± 6.1 | 0.315* |
| Lactate dehydrogenase (U/L) | 491 ± 338 | 556 ± 390 | 0.974** |
| Hemoglobin S (%) | 51.0 ± 22.4 | 47.6 ± 16.1 | 0.671* |
| Hemoglobin A (%) | 39.5 ± 23.1 | 39.7 ± 15.7 | 0.541* |
| Hemoglobin F (%) | 5.8 ± 6.7 | 2.5 ± 2.1 | 0.173** |
| **Pulmonary Artery Hypertension** | **4/10** | **9/11** | **0.049†** |
| **Intracardiac Shunt** | **1/10** | **7/11** | **0.012†** |

Values are mean ± SD
\* calculated based on one-way ANOVA
\*\* calculated based on non-parametric Mann-Whitney Test
† calculated based on $\chi^2$ -test

Obstruction in the small arteries in the lung causes increased pressure in the lung, leading to the complication known as pulmonary arterial hypertension (PAH). PAH is a debilitating disease even without SCD and puts more stress on the right side of the heart. We found that PAH was more common in the high 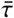 group ((Table 1), 4/10 vs. 9/11 (p=0.049, *χ*^2^-test). We also show that 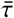 correlates with the pressure difference between the right ventricle and atrium of the heart. This difference can be calculated with the Bernoulli equation, Δ*P* = 4(TR_max_)^2^, where TR_max_ is the peak TRV and Δ*P* is the echocardiogram-estimated pulmonary artery pressure (PAP) (54). 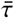 versus PAP is plotted in Fig. 6d (*p* = 0.031, linear regression). Medical guidance prescribes that PAH diagnosis should be confirmed with PAP using right heart catheterization (55), which is found to be well correlated with the echocardiogram-estimated PAP (56, 57). Furthermore, we found that intracardiac shunts are more common in the high 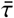 group (Table 1): 1/10 vs. 7/11 (*p* = 0.012, *χ*^2^-test). Intracardiac shunts are a form of a broader condition known as right-to-left shunting (RLS) where a portion of deoxygenated blood bypasses the lungs and returns to the systemic circulation without being oxygenated. RLS, which is increasingly being recognized in people with the HbSS sickle cell disease genotype (58, 59), may result in stroke. This is because it hinders resolution of small blood clots in the lungs by providing a passage for these clots to return to the circulation (60). Though etiology of RLS in SCD is unknown, the findings that high 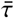 is associated with increased pulmonary pressure and intracardiac shunts are consistent with the hypothesis that deoxygenated HbS-containing RBCs obstruct the pulmonary vasculature and result in eventual formation of abnormal shunting (58).

Up to now we have discussed inflammation and hyperperfusion in the context of SCD and its relation to shear force. But there is another factor that influences the shear forces acting on sRBCs, namely whole blood viscosity. There are variety of features that determine blood viscosity: plasma viscosity, RBC deformability and aggregation, hematocrit (the fraction by volume of red blood cells in the blood) and the contribution of other vascular components—including leukocytes, endothelial cells and pericytes (61). In particular, reduced RBC deformability causes HbSS blood (the most common genotype in SCD) to be more viscous than normal blood (HbAA) at the same hematocrit level. This is not an issue for bulk flow in SCD patients, since the low hematocrit associated with severe anemia compensates to reduce the viscosity (62). However the microcirculation is potentially more problematic. In capillaries, where blood flow is maintained with low force, the viscosity is actually decreased further due to plasma skimming, approaching the viscosity of plasma (the Fåhræus–Lindqvist or sigma phenomenon). However, in the postcapillaries, it suddenly increases, the so-called “inversion phenomenon”. Notably, the hematocrit, which is exponentially related to blood viscosity, can increase up to 75% in this part of the microcirculation (63). The deformability of blood cells becomes a key factor to keep the flow resistance low in postcapillaries, so that RBC passage is completed before the rapid polymerization of the HbS after initial nucleation. Additionally, hemolysis increases the flow resistance by depleting nitric-oxide and activating the endothelium and leukocytes (63, 64). Abnormal cellular adhesion (along with other SCD features like increased RBC stiffness and hemolysis (63, 64)) can further contribute the flow resistance, increasing blood viscosity and therefore shear force in the post-capillary regions. Given the variation in *τ*(*f*) between high and low 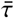 groups (as well as between catch and slip behaviors), the consequences of increased shear could vary widely from patient to patient. The same would be true when evaluating therapies that target reduction in the flow resistance by blocking cellular adhesive interactions (crizanlizumab, L-glutamine, purified poloxamer 188, etc.), and therapies that target improving the sickle RBC deformability (voxelotor). For example, crizanlizumab inhibits the adhesion of leukocytes, which would have a downstream effect of reduced flow resistance and reduced viscosity, thereby decreasing the shear force. This can be detrimental for a patient in the slip-like subgroup, due to increased adhesion lifetime at lower shear forces, defeating the purpose of the treatment. The results described in this section already highlight that the *τ*(*f*) profile for a given patient has clinical relevance. Further studies are needed to see if knowledge of this profile can be harnessed to tailor and improve the effectiveness of therapeutic interventions.

## CONCLUSIONS

In this study we examined sRBC-LN adhesion using a variety of complementary approaches. TIRM allowed us to visualize how the adhesion sites on the sRBC surface dynamically vary in both location and height in the moments before detachment. These complex dynamics are a requirement for the existence of catch bonding, which we subsequently observed in two different experiments that used increased shear force (either via sudden jumps or linear ramps) to detach sRBCs from an LN-functionalized surface. By extracting the mean bond lifetime versus force profiles for individual SCD patients using the experimental distributions of detachment events, we found that these profiles were remarkably heterogeneous among a cohort of 25 patients. Not only did the lifetime curves vary qualitatively (catch vs. slip), but also showed a wide range of magnitudes in the force regime typical of post-capillary vessels. Patients with large bond lifetimes at physiological forces were more likely to exhibit certain adverse SCD phenotypes: larger numbers of white blood cells (associated with inflammation), larger tricuspid regurgitation velocity (associated with hyperperfusion), pulmonary artery hypertension, and intracardiac shunting.

Our work is the first to observe dynamical heterogeneity in sRBC-LN adhesion and identify its clinical consequences. But there are still many open questions about these phenomena, at both molecular and phenotypic levels. In terms of molecular mechanisms, the biggest question is how the individual dynamics of protein-ligand complexes among the sRBC adhesion sites contribute to give an overall catch or slip lifetime behavior. Fig. 7 provides a schematic illustration of several possible explanations. Slip bonding is the simplest to rationalize, since it requires nothing more than individual complexes that form via single binding domains, which then become destabilized under increasing force. Catch bonding necessitates more elaborate mechanisms. Case I in Fig. 7 shows one possible route: if there are multiple available binding domains at the protein-ligand interface, remodeling of the interface under force might allow additional domains to engage, strengthening the bond (47, 65). But there are also alternatives, like case II in Fig. 7: here each protein-ligand complex is individually a slip bond, but there could be some coordination in the way these bonds are formed, either within a cluster at a given adhesion site, or among a group of sites (66, 67). And of course there could be even more complex mechanisms, involving mixtures of molecular catch and slip bonds determining the overall sRBC adhesion dynamics.

**Figure 7:**
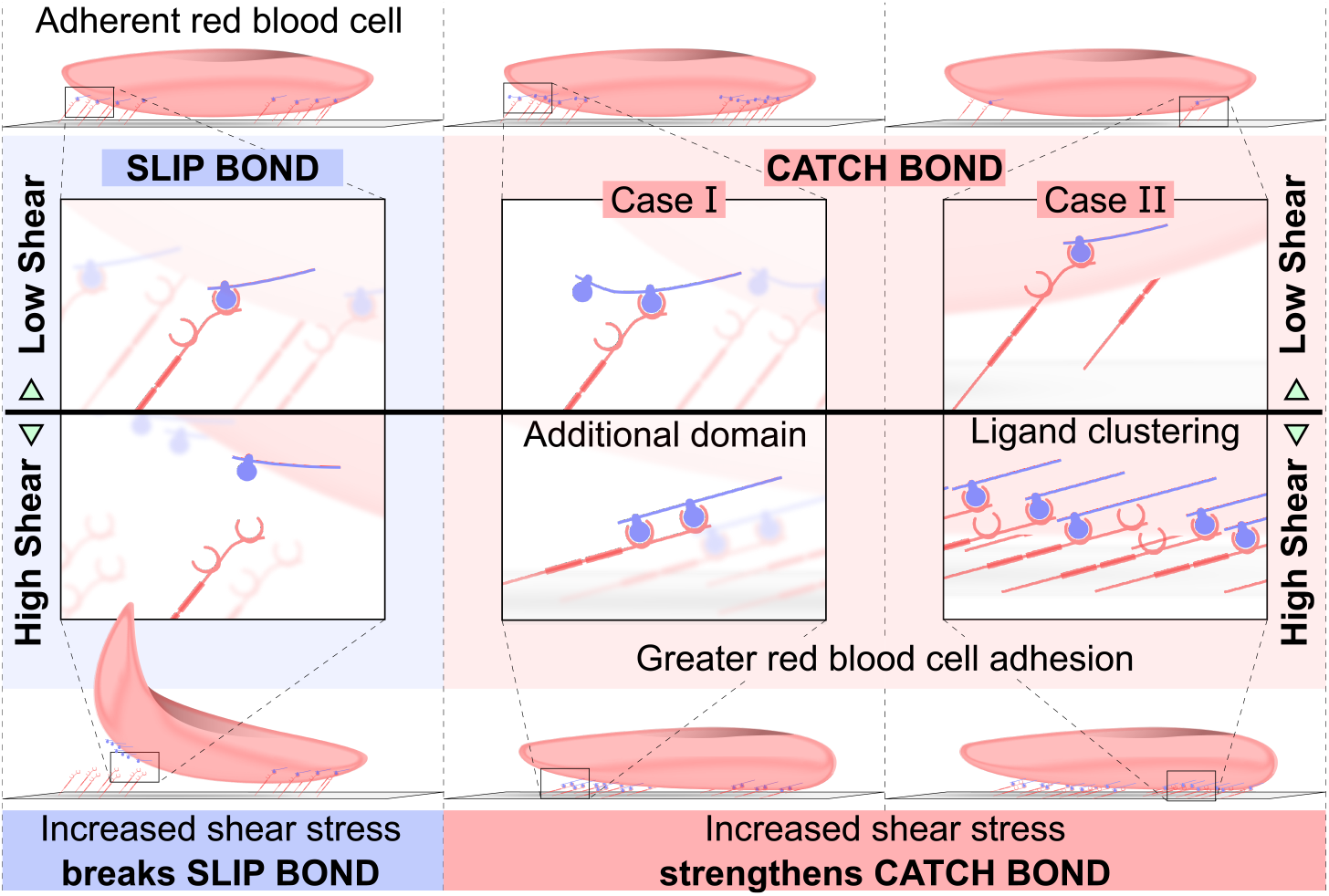
Possible molecular mechanisms underlying sRBC-LN slip and catch bond behaviors. Slip bonds could be mediated by individual protein-ligand complexes that have slip dynamics, for example binding via a single domain that becomes destabilized by increasing force. To get collective catch bond dynamics at the whole cell level, different scenarios are possible. For case I each individual protein-ligand complex is itself a catch bond, with additional binding domains that engage and strengthen the bond when force is applied to the complex. For case II each individual complex is a slip bond, but there could be coordination among clusters of the bonds that give rise to overall catch dynamics.

At the phenotypic level, it would be interesting to investigate how the dynamical heterogeneity of sRBC adhesion influences the progression of vaso-occlusive crises. One hypothesis is that when an occlusion at a single site is cleared, the resulting restoration of blood flow (reperfusion) causes increased blood velocities (hyperperfusion) in the downstream network of blood vessels. Does this lead to enhanced sRBC adhesion for patients with catch bonding, helping nucleate new occlusion sites and spreading the region of inflammation? How different is the probability of new occlusions and the consequent rate of spreading for patients with high versus low 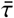? Can doing an *in vitro* measurement of sRBC adhesion dynamics for a patient allow us to predict the duration of a vaso-occlusive event? Answering these questions is especially important for assessing anti-adhesive therapies, including tinzaparin, IVIG, and sevuparin, whose primary outcome measures include time to resolution of acute vaso-occlusive complications in their clinical trials (68). Looking toward the future, it is clear that understanding the unique dynamical features of sRBC adhesion within a given patient could open up new avenues of personalized SCD therapies. But the usefulness of our approach is not limited to the context of SCD. The microfluidic experimental setup and data analysis techniques we describe here can be adapted to study the adhesion dynamics of various other cell types, i.e. leukocytes or cancer cells. We can also use our setup to quantify how adhesion is impacted by disease-related environmental conditions, for example oxygen deprivation (via an *in situ* gas exchanger (69)) or the presence of drugs.

## METHODS

### Blood sample collection and clinical study

All 33 subjects had phenotypic homozygous sickle hemoglobin (HbSS), and most had undergone echocardiography within a year of sample collection date. Clinical baseline implies without recent (within the last 2 weeks) history of VOC. Surplus EDTA-anticoagulated whole blood was obtained under an Institutional Review Board approved protocol and was tested for adhesion and Hb composition within 24 hours. Hb composition (fetal, HbF; and sickle, HbS, and adult, HbA (from transfusion)) was analyzed via High Performance Liquid Chromatography with a Bio-Rad Variant II Instrument (Bio-Rad, Montreal, QC, Canada) in the core clinical laboratory of University Hospitals Cleveland Medical Center (UHCMC). Clinical data were obtained from the Adult SCD Clinic at UHCMC, on dates contemporaneous with blood sample collection. Pulmonary arterial hypertension and intracardiac shunt diagnosis were obtained from the medical records.

### Microfluidic Adhesion Experiments

All microfluidic experiments are performed with the SCD Biochip.(8) Microchannels are coated with LN-1 for RBC-specific adhesion per protocol (8). LN-1 functionalized microfluidic channels are injected with 15 μl of unprocessed whole blood at shear forces within the range of post-capillary shear forces. In TIRM and ramped shear experiments, adherent RBCs are characterized after the non-adherent cells are removed by buffer wash at 0.5 dyne/cm^2^. To keep the shear force the same throughout the jumped shear experiments, non-adherent RBCs are kept continuously flowing at the predetermined shear forces during imaging. The net RBC adhesion was calculated with the following formula

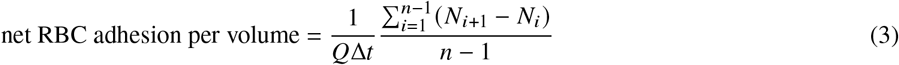

where *N_i_* is the number of adhered RBCs in a given image, *n* is the number of images captured at the flow rate *Q* and Δ*t* is the imaging interval. Adherent RBCs in the microchannels are visualized for quantification with Olympus CellSens software using an Olympus IX83 inverted microscope and QImaging EXi Blue CCD camera at 10x. Scanned images of jumped shear experiments covered actual area of 32.77 mm^2^ on the microchannels. Videos of ramped shear experiments covered an actual area of 0.34 mm^2^.

### Total internal reflection microscopy

TIRM is performed on an upright optical microscope (Olympus, BX51WI). In brief, a collimated laser beam coupled to a single-mode optical fiber (*λ* = 633 nm, Necsel Novatru) is directed onto the sides of a BK7 trapezoidal prism, allowing an angle of incidence *θ*_i_ = 68° (with respect to the normal of the top of the prism). The prism is held in place with a home-built mount that screws into the mechanical stage of the microscope. The sample is placed on top of the prism by putting refractive index matching oil in between. The total internal reflection condition is met between the bottom surface of the microfluidic cell (*n*_1_ = 1.51) and the aqueous medium of the sample (*n*_2_ = 1.33), as *θ*_i_ > *θ*_critical_, creating an evanescent wave that decays exponentially within a few hundred nanometers from the surface (*β*^-1^ ~ 100 nm), providing high sensitivity in the z-axis, normal to the surface. The laser power is adjusted between 18 – 20 mW using a computer control provided by the manufacturer. The video is collected from the top of the sample with a 40X objective (Olympus LUCPLFLN40X, working distance set to 2 mm), and then passed through a de-magnifying adapter (0.63 X) before the CCD camera (Hamamatsu ORCA-R2). The video of an individual sickle RBC detachment is acquired with a 25 ms integration time, at 25 – 28 frames per second and 2 × 2 pixel binning (maximum field of view = 344 *μ*m × 262 *μ*m). The bit depth of each pixel was set to 16 bits, and the file is saved in big tiff (.btf) format. The video is then cropped and saved as .mp4 in Matlab using an automated routine we developed.(38) The detachment is initiated in a flow cytometry staining buffer (RnD Systems, Minneapolis, MN) by ramped flow as described in Fig. 4A at a ramping rate of 2.5 dyne/cm^2^/min.

### Automated Processing of Ramped Shear Video Data

To automate and speed up the processing of image data from the ramped shear experiments, we developed a set of deep learning based object detection and tracking codes. The codes have been designed to set up a processing pipeline that can be used on a MATLAB platform. To start off the processing of microfluidic channel images generated in our whole blood detachment experiments, we need an image segmentation and object detection algorithm. To this end, we adapted the deep convolutional neural network developed in an earlier study from our group that can analyze bright field channel images from whole blood detachment experiments, and identify the adhered sRBCs (41). The tracking algorithm identifies an RBC as detached when the RBC leaves its initial adhered position. The algorithm does not analyze RBCs that entered the field of view after the ramping starts.

### Statistical Methods

Statistical analyses were performed using MATLAB. Data were tested for normality. Non-normally distributed data were analyzed using a non-parametric Mann-Whitney U-test for two independent groups. Normally distributed data were analyzed using a one-way ANOVA test for two independent groups. The chi-square method was utilized to test the associations between two categorical variables. *p* < 0.05 was chosen to indicate a significant difference.

## ACKNOWLEDGEMENTS

This work was supported by National Heart Lung and Blood Institute R01HL133574, OT2HL152643, T32HL134622, K25HL159358, National Heart Lung and Blood and the National Center for Complementary & Integrative Health (NCCIH) U54HL143541, and National Science Foundation CAREER Awards 1552782 and 1651560. This work was partially supported by the National Science Foundation CAREER Award, NSF No. 2023525. The authors acknowledge with gratitude the contributions of research participants and clinicians at Seidman Cancer Center (University Hospitals, Cleveland).

## AUTHORSHIP CONTRIBUTIONS

U. G, S. I., M. H. and U. A. G., developed the idea and designed the study. U. G., S. D., Y. M. and R. A. performed the experiments. U. G., S. I., G. S., S. D., Y.M., A. B. and R.A. analyzed the data. U. G., S. I., G. S., A. B., J. A. L., C. L. W., M. H. and U.A.G. discussed and interpreted the data. S. I. and G. S. performed the modelling. U. G., S. I., and M. H. wrote the manuscript, U.G. and S. I. prepared the figures and table. J. A. L., C. L. W., M. H. and U.A.G., edited the manuscript. U. G. and S.I. contributed equally.

## SUPPLEMENTARY INFORMATION

### Shear force and wall shear stress calculations

Shear forces acting on the sickle red blood cells (sRBC) are calculated based on the assumption of average homozygous sickle cell blood viscosity is 3.73 cP as measured previously within the same microfluidic channels (62). Viscosity of the buffer solution is assumed identical to water. Shear force (*F_s_*) acting on the RBCs is obtained by the following solutions of Stokes flow for spheroid particles (70):

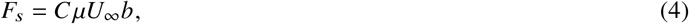

where *C* is constant, *μ* is the dynamic viscosity of the fluid, 2*b* is the length along the minor axis, 2*a* is the length along the major axis of the RBC. With the free stream velocity (*U*_∞_) tangential (along the major axis of the RBC), *C* can be calculated by the following equation:

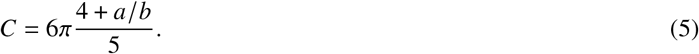

Wall shear stress on the adhesion surface was calculated using the following equation:

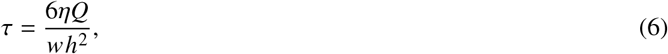

where *η* is the kinematic viscosity, *Q* is the flow rate, *w* is the width and *h* is the height of the microchannel. The physiological range of wall shear stress in post-capillary venules ranging in size (*d*) from 10-50 *μ*m was estimated based on the following empirical formula derived by Koutsiaris *et al*. (71):

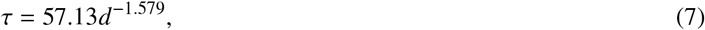

where *τ* is in pascals and *d* is in micrometers.

### Maximum likelihood estimation for bond lifetime model parameters

To fit our sRBC-LN binding model to the data from our detachment experiments, we utilized a maximum likelihood estimation (MLE) scheme. The fitting procedure is outlined here. We start by establishing some relevant nomenclature. Let us denote our model by 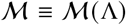 where Λ = {*k*_1_, *k*_2_, *d*_1_, *d*_2_} is the set of free parameters governing the model. Each detachment experiment on a certain sample is performed at a specific force ramp *r_i_*, and the ramps span a set *i* = 1, 2,…*N_r_*. The data consists of observed individual cellular detachment events under shear force pulling. We bin the observed force spectrum into *N_w_* bins of width *w* to construct force based histograms. Let the number of individual experimental detachment events observed for the *j*-th force bin centred at *f_j_* (i.e. for *f_j_* – *w*/2 < *f* < *f_j_* + *w*/2) be *n_j_*. The data for a certain sample for the *i*-th force ramp can then be cast in the form 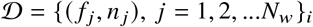. We now need a map between 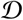 and 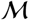. Given the model 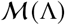, the conditional probability of obtaining a single detachment event in the *j*-th force bin for an experiment performed at the z-th force ramp *r_i_* can be constructed using:

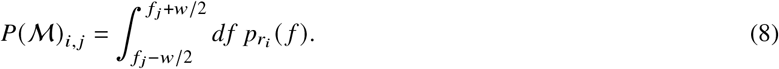

*p_r_*(*f*) is the underlying probability distribution of rupture forces governing an experiment performed at force ramp *r*. The overall likelihood of observing the full data set of (independent) outcomes 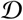 from *N_r_* force ramped experiments on a certain patient sample, given a particular model 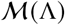, can be obtained from:

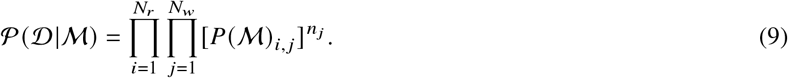

The rupture force probability distribution can be directly related to the survival probability distribution Σ*_r_*(*f*):

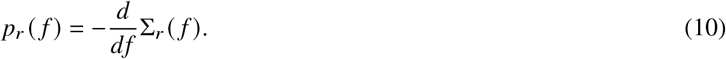

Since we already have a postulated analytical form for Σ*_r_*(*f*) from Eqs. (1) and (2) in the main text, we can compute the model-dependent functional forms of *p_r_*(*f*), 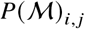 and 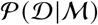. Model fitting now becomes a straightforward problem of obtaining estimates for the model parameters {*k*_1_, *k*_2_, *d*_1_, *d*_2_} that maximize the experimentally observed likelihood function 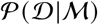 given its functional form. We can utilize this fitting scheme to independently fit data from the detachment experiments for each individual patient, yielding a different set of parameters for each patient.

### Clinical phenotypes of the catch-like versus slip-like patient subgroups

The comparison of clinical data from complete blood counts between the catch-like and slip-like patient subgroups (Supplementary Table 1) did not indicate a strong statistical difference.

**Supplementary Table 1:** Clinical phenotype of the sickle cell disease patients with slip-like and catch-like adhesion.

| Parameters (Units) | Subjects with<br>slip-like adhesion<br>N=16 | Subjects with<br>catch-like adhesion<br>N=9 | p |
| --- | --- | --- | --- |
| Age | 31.5 ± 7.0 | 36.1 ± 9.1 | 0.169* |
| White Blood Cells (10<sup>9</sup>/L) | 11.9 ± 3.2 | 12.8 ± 2.0 | 0.465* |
| Absolute Neutrophil Count (10<sup>6</sup>/L) | 7539 ± 2876 | 7776 ± 2102 | 0.841* |
| Platelets (10<sup>9</sup>/L) | 361 ± 68 | 364 ± 149 | 0.952* |
| Mean Corpuscular Volume (fL) | 90.0 ± 5.0 | 84.1 ± 17.6 | 0.974** |
| Hemoglobin (g/dL) | 8.4 ± 1.6 | 9.0 ± 1.7 | 0.405* |
| Hematocrit (%) | 24.8 ± 3.6 | 25.9 ± 4.5 | 0.532* |
| Tricuspid Regurgitation Velocity, Peak (m/s) | 2.73 ± 0.94 | 2.87 ± 0.63 | 0.386** |
| Red Blood Cells (10<sup>12</sup>/L) | 2.8 ± 0.4 | 2.9 ± 0.6 | 0.499* |
| Absolute Reticulocyte Count (10<sup>9</sup>/L) | 462 ± 208 | 434 ± 121 | 0.720* |
| Reticulocyte percent (%) | 16.6 ± 6.4 | 15.8 ± 5.2 | 0.780* |
| Lactate dehydrogenase (U/L) | 591 ± 422 | 405 ± 168 | 0.275** |
| Hemoglobin S (%) | 47.4 ± 19.3 | 52.1 ± 18.8 | 0.563* |
| Hemoglobin A (%) | 39.5 ± 19.3 | 34.0 ± 19.6 | 0.513* |
| Hemoglobin F (%) | 3.9 ± 5.3 | 4.1 ± 4.7 | 0.834** |

Values are Mean ± SD
\* calculated based on one-way ANOVA
\*\* calculated based on non-parametric Mann-Whitney Test

**Supplementary Figure 1:**
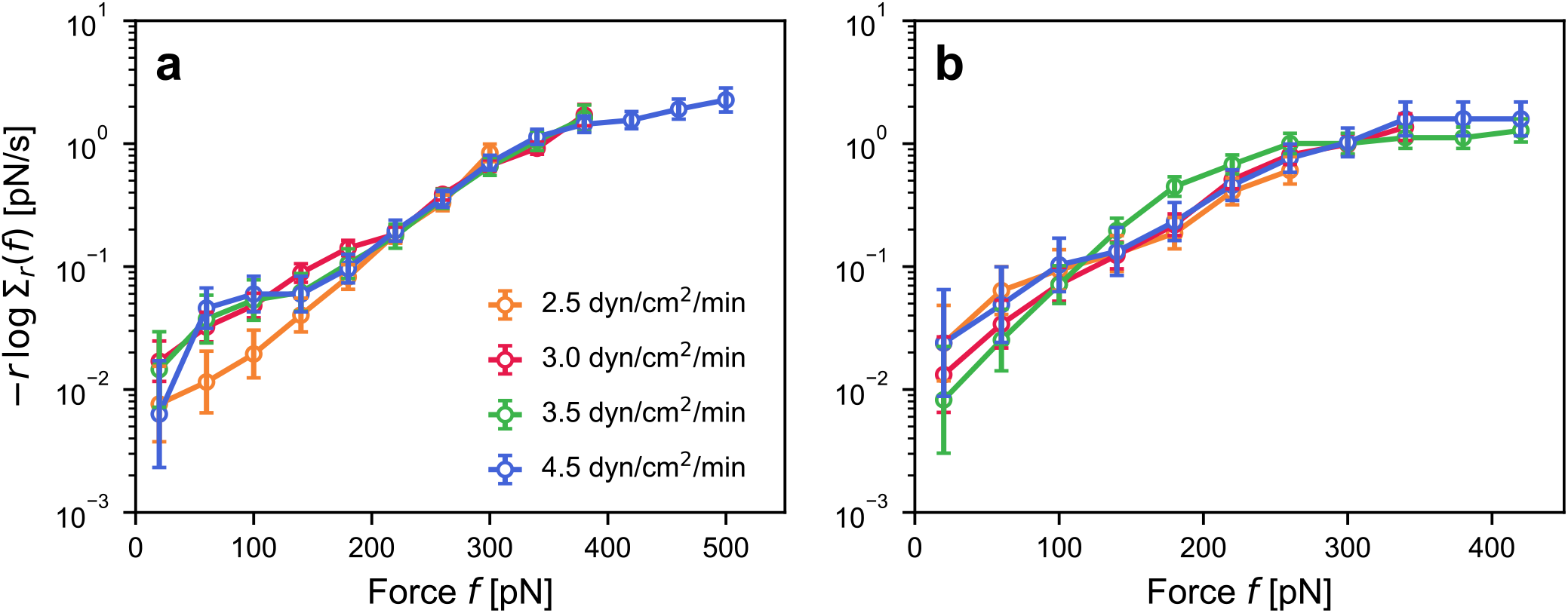
**(a,b)** Plots of −*r* log Σ_*r*_(*f*) vs. force *f* for two different patients, using the ramp rates *r* shown in main text Fig. 4. The fact that the curves at different ramp rates are similar for each patient indicates that the sRBC-LN bond is in the pure adiabatic regime. In contrast, for a heterogeneous system the curves would clearly diverge from one another at larger *f*, as seen in the examples analyzed in Ref. (42).

**Supplementary Figure 2:**
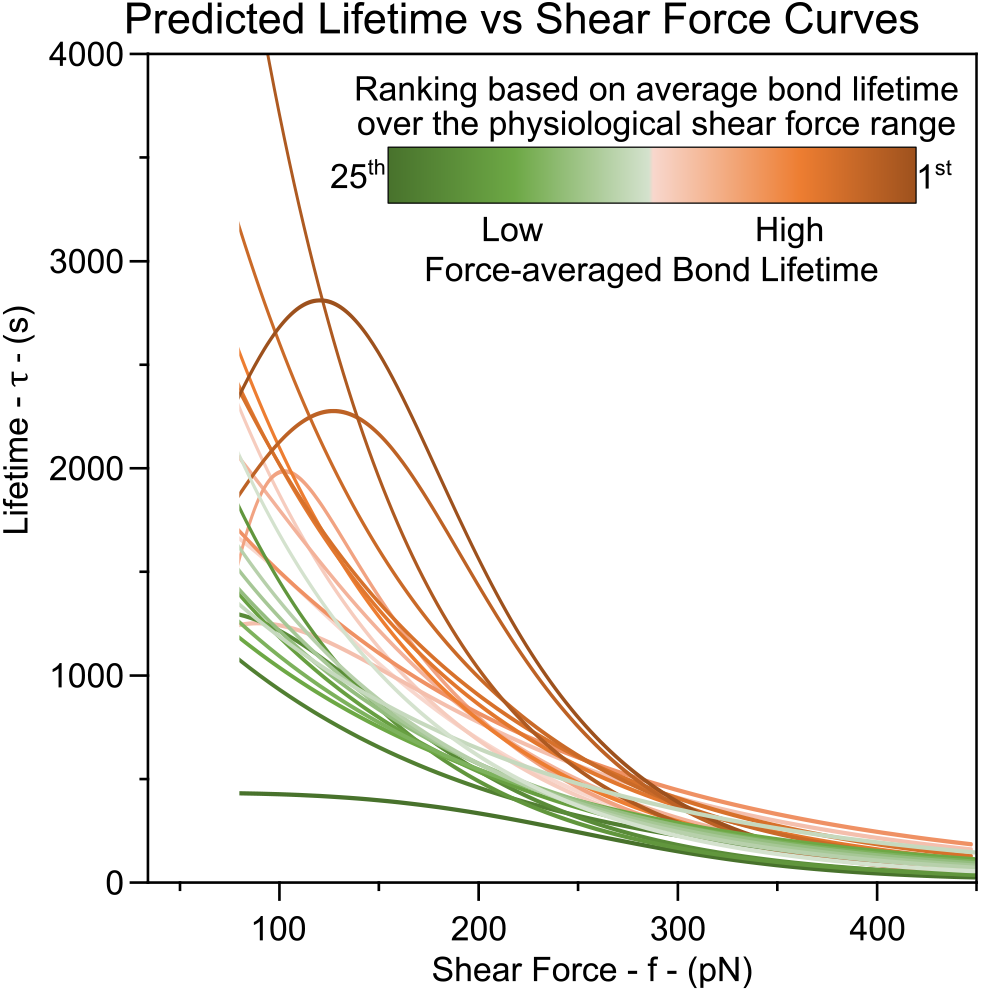
Patient groups based on the force-averaged bond lifetime 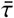, annotated by color. Patients are split into two groups (low & high force-averaged bond lifetime) from the median of their 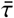 values.

## REFERENCES

1. Abdu, A., J. Gómez-Márquez, and T. K. Aldrich, 2008. The oxygen affinity of sickle hemoglobin. Respiratory Physiology & Neurobiology 161:92–94.

2. Noomuna, P., M. Risinger, S. Zhou, K. Seu, Y. Man, R. An, D. A. Sheik, J. Wan, J. A. Little, U. A. Gurkan, et al., 2020. Inhibition of Band 3 tyrosine phosphorylation: a new mechanism for treatment of sickle cell disease. British Journal of Haematology 190:599–609.

3. Frenette, P. S., G. F. Atweh, et al., 2007. Sickle cell disease: old discoveries, new concepts, and future promise. The Journal of Clinical Investigation 117:850–858.

4. Man, Y., E. Kucukal, R. An, Q. D. Watson, J. Bosch, P. A. Zimmerman, J. A. Little, and U. A. Gurkan, 2020. Microfluidic assessment of red blood cell mediated microvascular occlusion. Lab on a Chip 20:2086–2099.

5. Inanc, M. T., I. Demirkan, C. Ceylan, A. Ozkan, O. Gundogdu, U. Goreke, U. A. Gurkan, and M. B. Unlu, 2021. Quantifying the influences of radiation therapy on deformability of human red blood cells by dual-beam optical tweezers. RSC Advances 11:15519–15527.

6. Man, Y., R. An, K. Monchamp, Z. Sekyonda, E. Kucukal, C. Federici, W. J. Wulftange, U. Goreke, A. Bode, V. Sheehan, et al. OcclusionChip: a functional microcapillary occlusion assay complementary to ektacytometry for detection of small-fraction red blood cells with abnormal deformability. Frontiers in Physiology 1526.

7. Goreke, U., A. Bode, S. Yaman, U. A. Gurkan, and N. G. Durmus, 2022. Size and density measurements of single sickle red blood cells using microfluidic magnetic levitation. Lab Chip 22:683–696.

8. Alapan, Y., C. Kim, A. Adhikari, K. E. Gray, E. Gurkan-Cavusoglu, J. A. Little, and U. A. Gurkan, 2016. Sickle cell disease biochip: a functional red blood cell adhesion assay for monitoring sickle cell disease. Translational Research 173:74–91.

9. An, R., Y. Man, K. Cheng, T. Zhang, C. Chen, E. Kucukal, W. Wulftange, U. Goreke, A. Bode, L. Nayak, et al., 2022. Sickle red blood cell derived extracellular vesicles activate endothelial cells and enhance sickle red cell adhesion mediated by von Willebrand factor. bioRxiv.

10. Man, Y., E. Kucukal, R. An, A. Bode, J. A. Little, and U. A. Gurkan, 2021. Standardized microfluidic assessment of red blood cell–mediated microcapillary occlusion: Association with clinical phenotype and hydroxyurea responsiveness in sickle cell disease. Microcirculation 28:e12662.

11. Platt, O. S., 2000. Sickle cell anemia as an inflammatory disease. Journal of Clinical Investigation 106:337–338.

12. Man, Y., U. Goreke, E. Kucukal, A. Hill, R. An, S. Liu, A. Bode, A. Solis-Fuentes, L. V. Nayak, J. A. Little, et al., 2020. Leukocyte adhesion to P-selectin and the inhibitory role of Crizanlizumab in sickle cell disease: A standardized microfluidic assessment. Blood Cells, Molecules, and Diseases 83:102424.

13. Hines, P. C., Q. Zen, S. N. Burney, D. A. Shea, K. I. Ataga, E. P. Orringer, M. J. Telen, and L. V. Parise, 2003. Novel epinephrine and cyclic AMP-mediated activation of BCAM/Lu-dependent sickle (SS) RBC adhesion. Blood, The Journal of the American Society of Hematology 101:3281–3287.

14. Solovey, A., Y. Lin, P. Browne, S. Choong, E. Wayner, and R. P. Hebbel, 1997. Circulating activated endothelial cells in sickle cell anemia. New England Journal of Medicine 337:1584–1590.

15. Hillery, C. A., M. C. Du, W. C. Wang, and J. P. Scott, 2000. Hydroxyurea therapy decreases the in vitro adhesion of sickle erythrocytes to thrombospondin and laminin. British Journal of Haematology 109:322–327.

16. Bell, G., 1978. Models for the specific adhesion of cells to cells. Science 200:618–27.

17. Dembo, M., D. Torney, K. Saxman, and D. Hammer, 1988. The reaction-limited kinetics of membrane-to-surface adhesion and detachment. Proceedings of the Royal Society of London. Series B. Biological Sciences 234:55–83.

18. Marshall, B., M. Long, J. Piper, T. Yago, R. McEver, and C. Zhu, 2003. Direct observation of catch bonds involving cell-adhesion molecules. Nature 423:190–193.

19. Thomas, W. E., E. Trintchina, M. Forero, V. Vogel, and E. V. Sokurenko, 2002. Bacterial adhesion to target cells enhanced by shear force. Cell 109:913–923.

20. Chakrabarti, S., M. Hinczewski, and D. Thirumalai, 2017. Phenomenological and microscopic theories for catch bonds. Journal of Structural Biology 197:50–56.

21. Zhuravlev, P. I., M. Hinczewski, S. Chakrabarti, S. Marqusee, and D. Thirumalai, 2016. Force-dependent switch in protein unfolding pathways and transition-state movements. Proceedings of the National Academy of Sciences 113:E715–E724.

22. Rieger, H., H. Y. Yoshikawa, K. Quadt, M. A. Nielsen, C. P. Sanchez, A. Salanti, M. Tanaka, and M. Lanzer, 2015. Cytoadhesion of Plasmodium falciparum–infected erythrocytes to chondroitin-4-sulfate is cooperative and shear enhanced. Blood 125:383–391.

23. Kikkawa, Y., and J. H. Miner, 2005. Lutheran/B-CAM: a laminin receptor on red blood cells and in various tissues. Connective Tissue Research 46:193–199.

24. Murphy, M. M., M. A. Zayed, A. Evans, C. E. Parker, K. I. Ataga, M. J. Telen, and L. V. Parise, 2005. Role of Rap1 in promoting sickle red blood cell adhesion to laminin via BCAM/LU. Blood 105:3322–3329.

25. Udani, M., Q. Zen, M. Cottman, N. Leonard, S. Jefferson, C. Daymont, G. Truskey, and M. J. Telen, 1998. Basal cell adhesion molecule/lutheran protein. The receptor critical for sickle cell adhesion to laminin. The Journal of Clinical Investigation 101:2550–2558.

26. Deng, Y., D. P. Papageorgiou, H.-Y. Chang, S. Z. Abidi, X. Li, M. Dao, and G. E. Karniadakis, 2019. Quantifying shear-induced deformation and detachment of individual adherent sickle red blood cells. Biophysical Journal 116:360–371.

27. Zen, Q., M. Batchvarova, C. A. Twyman, C. E. Eyler, H. Qiu, L. M. De Castro, and M. J. Telen, 2004. B-CAM/LU expression and the role of B-CAM/LU activation in binding of low-and high-density red cells to laminin in sickle cell disease. American Journal of Hematology 75:63–72.

28. Kucukal, E., J. A. Little, and U. A. Gurkan, 2018. Shear dependent red blood cell adhesion in microscale flow. Integrative Biology 10:194–206.

29. Kato, G. J., F. B. Piel, C. D. Reid, M. H. Gaston, K. Ohene-Frempong, L. Krishnamurti, W. R. Smith, J. A. Panepinto, D. J. Weatherall, F. F. Costa, et al., 2018. Sickle cell disease. Nature Reviews Disease Primers 4:1–22.

30. Yuan, C., E. Quinn, E. Kucukal, S. Kapoor, U. A. Gurkan, and J. A. Little, 2019. Priapism, hemoglobin desaturation, and red blood cell adhesion in men with sickle cell anemia. Blood Cells, Molecules, and Diseases 79:102350.

31. Alapan, Y., J. A. Little, and U. A. Gurkan, 2014. Heterogeneous red blood cell adhesion and deformability in sickle cell disease. Scientific Reports 4.

32. An, R., Y. Man, S. Iram, E. Kucukal, M. N. Hasan, Y. Huang, U. Goreke, A. Bode, A. Hill, K. Cheng, et al., 2021. Point-of-Care microchip electrophoresis for integrated anemia and hemoglobin variant testing. Lab on a Chip.

33. Vekilov, P. G., 2007. Sickle-cell haemoglobin polymerization: is it the primary pathogenic event of sickle-cell anaemia? British Journal of Haematology 139:173–184.

34. Zhou, Z., P. Thiagarajan, M. M. Udden, J. A. López, and P. Guchhait, 2011. Erythrocyte membrane sulfatide plays a crucial role in the adhesion of sickle erythrocytes to endothelium. Thrombosis and Haemostasis 105:1046–1052.

35. Guchhait, P., P. Thiagarajan, and J. A. Lopez, 2007. Cell Membrane Sulfatide Promotes Sickle Cell Adhesion to Endothelium.

36. Xu, X., A. K. Efremov, A. Li, L. Lai, M. Dao, C. T. Lim, and J. Cao, 2013. Probing the cytoadherence of malaria infected red blood cells under flow. PloS One 8:e64763.

37. Doicu, A., A. A. Vasilyeva, D. S. Efremenko, C. L. Wirth, and T. Wriedt, 2019. A light scattering model for total internal reflection microscopy of geometrically anisotropic particles. Journal of Modern Optics 66:1139–1151.

38. Rashidi, A., S. Domínguez-Medina, J. Yan, D. S. Efremenko, A. A. Vasilyeva, A. Doicu, T. Wriedt, and C. L. Wirth, 2020. Developing scattering morphology resolved total internal reflection microscopy (SMR-TIRM) for orientation detection of colloidal ellipsoids. Langmuir 36:13041–13050.

39. Maciaszek, J. L., B. Andemariam, K. Abiraman, and G. Lykotrafitis, 2014. AKAP-dependent modulation of BCAM/Lu adhesion on normal and sickle cell disease RBCs revealed by force nanoscopy. Biophysical Journal 106:1258–1267.

40. Rupprecht, P., L. Golé, J.-P. Rieu, C. Vézy, R. Ferrigno, H. C. Mertani, and C. Riviere, 2012. A tapered channel microfluidic device for comprehensive cell adhesion analysis, using measurements of detachment kinetics and shear stress-dependent motion. Biomicrofluidics 6:014107.

41. Praljak, N., S. Iram, U. Goreke, G. Singh, A. Hill, U. A. Gurkan, and M. Hinczewski, 2021. Integrating deep learning with microfluidics for biophysical classification of sickle red blood cells adhered to laminin. PLoS Computational Biology 17:e1008946.

42. Hinczewski, M., C. Hyeon, and D. Thirumalai, 2016. Directly measuring single-molecule heterogeneity using force spectroscopy. Proceedings of the National Academy of Sciences 113:E3852–E3861.

43. Barsegov, V., and D. Thirumalai, 2005. Dynamics of unbinding of cell adhesion molecules: transition from catch to slip bonds. Proceedings of the National Academy of Sciences 102:1835–1839.

44. Dudko, O. K., G. Hummer, and A. Szabo, 2008. Theory, analysis, and interpretation of single-molecule force spectroscopy experiments. Proceedings of the National Academy of Sciences 105:15755–15760.

45. Evans, E., A. Leung, V. Heinrich, and C. Zhu, 2004. Mechanical switching and coupling between two dissociation pathways in a P-selectin adhesion bond. Proceedings of the National Academy of Sciences 101:11281–11286.

46. Elms, P. J., J. D. Chodera, C. Bustamante, and S. Marqusee, 2012. The molten globule state is unusually deformable under mechanical force. Proceedings of the National Academy of Sciences 109:3796–3801.

47. Chakrabarti, S., M. Hinczewski, and D. Thirumalai, 2014. Plasticity of hydrogen bond networks regulates mechanochemistry of cell adhesion complexes. Proceedings of the National Academy of Sciences 111:9048–9053.

48. Adhikari, S., J. Moran, C. Weddle, and M. Hinczewski, 2018. Unraveling the mechanism of the cadherin-catenin-actin catch bond. PLoS computational biology 14:e1006399.

49. Thomas, W., 2008. Catch bonds in adhesion. Annual Reviews of Biomedical Engineering 10:39–57.

50. Beste, M. T., and D. A. Hammer, 2008. Selectin catch–slip kinetics encode shear threshold adhesive behavior of rolling leukocytes. Proceedings of the National Academy of Sciences 105:20716–20721.

51. Nath, K. A., and R. P. Hebbel, 2015. Sickle cell disease: renal manifestations and mechanisms. Nature Reviews Nephrology 11:161–171.

52. Adams, R., V. McKie, F. Nichols, E. Carl, D.-L. Zhang, K. McKie, R. Figueroa, M. Litaker, W. Thompson, and D. Hess, 1992. The use of transcranial ultrasonography to predict stroke in sickle cell disease. New England Journal of Medicine 326:605–610.

53. Hebbel, R. P., J. D. Belcher, G. M. Vercellotti, et al., 2020. The multifaceted role of ischemia/reperfusion in sickle cell anemia. The Journal of Clinical Investigation 130:1062–1072.

54. Parasuraman, S., S. Walker, B. L. Loudon, N. D. Gollop, A. M. Wilson, C. Lowery, and M. P. Frenneaux, 2016. Assessment of pulmonary artery pressure by echocardiography—a comprehensive review. IJC Heart & Vasculature 12:45–51.

55. McLaughlin, V. V., S. L. Archer, D. B. Badesch, R. J. Barst, H. W. Farber, J. R. Lindner, M. A. Mathier, M. D. McGoon, M. H. Park, R. S. Rosenson, et al., 2009. ACCF/AHA 2009 expert consensus document on pulmonary hypertension: a report of the American College of Cardiology Foundation Task Force on expert consensus documents and the American Heart Association developed in collaboration with the American College of Chest Physicians; American Thoracic Society, Inc.; and the Pulmonary Hypertension Association. Journal of the American College of Cardiology 53:1573–1619.

56. Currie, P. J., J. B. Seward, K.-L. Chan, D. A. Fyfe, D. J. Hagler, D. D. Mair, G. S. Reeder, R. A. Nishimura, and A. J. Tajik, 1985. Continuous wave Doppler determination of right ventricular pressure: a simultaneous Doppler-catheterization study in 127 patients. Journal of the American College of Cardiology 6:750–756.

57. Yock, P. G., and R. L. Popp, 1984. Noninvasive estimation of right ventricular systolic pressure by Doppler ultrasound in patients with tricuspid regurgitation. Circulation 70:657–662.

58. Hambley, B. C., R. A. Rahman, M. Reback, M. A. O’riordan, N. Langer, R. C. Gilkeson, M. Ginwalla, J. A. Little, and R. Schilz, 2019. Intracardiac or intrapulmonary shunts were present in at least 35% of adults with homozygous sickle cell disease followed in an outpatient clinic. Haematologica 104:e1.

59. Kucukal, E., Y. Man, E. Quinn, N. Tewari, R. An, A. Ilich, N. S. Key, J. A. Little, and U. A. Gurkan, 2020. Red blood cell adhesion to ICAM-1 is mediated by fibrinogen and is associated with right-to-left shunts in sickle cell disease. Blood advances 4:3688–3698.

60. Abushora, M. Y., N. Bhatia, Z. Alnabki, M. Shenoy, M. Alshaher, and M. F. Stoddard, 2013. Intrapulmonary shunt is a potentially unrecognized cause of ischemic stroke and transient ischemic attack. Journal of the American Society of Echocardiography 26:683–690.

61. Baskurt, O. K., and H. J. Meiselman, 2003. Blood Rheology and Hemodynamics. Seminars in Thrombosis and Hemostasis 29:435–450.

62. Kucukal, E., Y. Man, A. Hill, S. Liu, A. Bode, R. An, J. Kadambi, J. A. Little, and U. A. Gurkan, 2020. Whole blood viscosity and red blood cell adhesion: Potential biomarkers for targeted and curative therapies in sickle cell disease. American Journal of Hematology 95:1246–1256.

63. Klug, P. P., L. S. Lessin, and P. Radice, 1974. Rheological aspects of sickle cell disease. Archives of Internal Medicine 133:577–590.

64. Nader, E., S. Skinner, M. Romana, R. Fort, N. Lemonne, N. Guillot, A. Gauthier, S. Antoine-Jonville, C. Renoux, M.-D. Hardy-Dessources, et al., 2019. Blood rheology: key parameters, impact on blood flow, role in sickle cell disease and effects of exercise. Frontiers in Physiology 10:1329.

65. Springer, T. A., 2009. Structural basis for selectin mechanochemistry. Proceedings of the National Academy of Sciences 106:91–96.

66. Björnham, O., and O. Axner, 2010. Catch-bond behavior of bacteria binding by slip bonds. Biophysical Journal 99:1331–1341.

67. Lin, J., Y. Wang, and J. Qian, 2021. Effects of domain unfolding and catch-like dissociation on the collective behavior of integrin–fibronectin bond clusters. Acta Mechanica Sinica 37:229–243.

68. Moerdler, S., and D. Manwani, 2018. New insights into the pathophysiology and development of novel therapies for sickle cell disease. Hematology 2014, the American Society of Hematology Education Program Book 2018:493–506.

69. Kim, M., Y. Alapan, A. Adhikari, J. A. Little, and U. A. Gurkan, 2017. Hypoxia-enhanced adhesion of red blood cells in microscale flow. Microcirculation 24:e12374.

70. Johnson, R. W., 2016. Handbook of Fluid Dynamics. Crc Press.

71. Koutsiaris, A. G., S. V. Tachmitzi, N. Batis, M. G. Kotoula, C. H. Karabatsas, E. Tsironi, and D. Z. Chatzoulis, 2007. Volume flow and wall shear stress quantification in the human conjunctival capillaries and post-capillary venules in vivo. Biorheology 44:375–386.

